# Effect of Application of Titanium Dioxide in the Management of Fusarium Wilt and Fruit Yield of Some Tomato Accessions

**DOI:** 10.1101/2022.09.02.506275

**Authors:** R. O. Olanrewaju, A. R. Popoola, C. G. Afolabi, J. G. Bodunde, S. A. Ganiyu

## Abstract

Tomato (*Solanum lycopoersicum* L.) is often threatened by wilt disease caused by *Fusarium oxysporium* f.sp *lycopersici*. Titanium dioxide (TiO_2_) has been reported to promote plant growth and reduce disease severity. This experiment was carried out to investigate the effects of TiO_2_ application on incidence and severity of *Fusarium* wilt as well as yield indices of three susceptible tomato accessions. A 3 × 5 factorial experiment fitted into Completely Randomized Design and Randomized Complete Block Design in both the screenhouse and the field, respectively. All experiments were set up with three replications. The treatments consisted of three tomato accessions (CPTTO/19/191, CPTTO/19/193 and CPTTO/19/195) and TiO_2_ with four concentrations (0.3, 0.7, 1.0 and 1.3 ml/l) was applied using soil drenching. Plots without TiO_2_ application served as the control. In both screenhouse and the field experiments, application of 1.3 ml/l TiO_2_ significantly reduced the incidence and severity of *Fusarium wilt* with better yield of tomato fruit in the three accessions than the control plots and pots. The study concluded that application of TiO_2_ at 1.3 ml/l reduced incidence and severity of *Fusaruim* wilt of tomato and increased the yield of tomato.

## INTRODUCTION

Tomato (*Solanum lycopersicum* L.) was native to tropical America, but yet grown all over the world (Arah *et al*., 2015). Tomatoes production accounts for about 4.8 million hectares of harvested land area globally with an estimated production of 16.2 million tonnes (FAOSTAT, 2014). China leads world tomato production with about 50 million tonnes followed by India with 17.5 million tonnes (FAOSTAT, 2014). In Africa, total tomato production for 2012 was 17.938 million tons with Egypt leading the continent with 8.625 million tonnes, followed by Nigeria with 1.56 million tonnes (Arah *et al*., 2015).

Tomato production can serve as a source of income for most rural and peri-urban producers in most developing countries. The tomato industry has been identified as an area that has the ability for poverty reduction because of its potential for growth and employment creation (Anang *et al*., 2013). Tomato has become an important cash and industrial crop in many parts of the world (Ayandiji *et al*., 2011) not only because of its economic importance but also its nutritional value to human diet and subsequent importance in human health (Willcox *et al*., 2003). In Nigeria, production of tomato has improved the livelihood of most rural and peri-urban farmers (Adenuga *et al*., 2013).

Some of the common problems of tomato production are pest and diseases. Diseases which include *Fusarium* wilt, Bacterial wilt, Anthracnose, Verticillium wilt etc (Ebimieowei *et al*., 2013). These pathogens low quality and insufficient quantity of tomato (Robinson and Kolavalli, 2010). *Fusarium oxysporum* f.sp. *lycopersici* (Sacc.) Snyder and Hans, is a soil borne plant pathogen that causes *Fusarium* wilt specifically on tomato (Rai *et al*., 2011). *Fusarium oxysporum* f. sp. *lycopersici* has become one of the most damaging pathogens wherever tomatoes are grown intensively because it grows endophytically and persists in infested soils (Agrios, 1997). *Fusarium oxysporum* f.sp. *lycopersici* is a known pathogen of tomato plant which is an economically important crop (Suarez-Estrella *et al*., 2007). Tomato yield is significantly reduced by the pathogen. Healthy plants can become infected if the soil in which they are growing is infested with the pathogen (Agrios, 1988). Once an area becomes infected with *F. oxysporum*, it usually remains so indefinitely (Agrios, 1997). The pathogen invades the vascular tissues, grows in the vascular bundles and inhibits water flow consequently causing wilting and ultimately leading to death of the plant (Davies, 1982). *Fusarium* wilt leads to an average yield loss of 50 % in tomato, reduces farmer’s income and family intake of vitamin A (Ajigbola *et al*., 2013). It constitutes serious threat to food security in Sub-Saharan Africa, especially in the coastal regions (Popoola *et al*., 2012).

Frequent use of synthetic fungicides to tackle this lethal disease of tomato has often resulted in environmental damage and increases pathogen resistance (Ogzonen *et al*., 2001). Resistance of *F. oxysporum* f. sp. *lycopersici* to synthetic fungicides necessitates the use of alternative control method to wilt caused by the pathogen (Ogzonen *et al*., 2001). Application of TiO_2_ has been found to show an excellent efficacy in rice (*Oryza sativa* L) and maize (*Zea Mays* L) by reducing the effect of *Curvularia* leaf spot and bacterial leaf blight disease incidence and severity (Chao *et al*. 2005; Hamzat etal., 2022). Hence, the objective of this work was to examine the effect of application of Titanium dioxide in the management of *Fusarium* wilt and fruit yield of three tomato accessions.

## MATERIALS AND METHODS

### Experimental site, designs and treatments

The experiment was carried out at Federal University of Agriculture Abeokuta (FUNAAB) DelPHE-5 Research farm, Ogun State, Nigeria. The location enjoys tropical climate with uni-modal peak rainfall between June and November, average annual and monthly rainfall of 1, 220 mm and 102 mm, respectively, as well as monthly maximum and minimum temperature ranges of 29–36 and 22–35°C, respectively (Kilanko-Oluwasanya, 2009). Humidity is lowest (37%– 54%) at the peak of dry season in February and highest at the peak of the rainy season between June and September (78%–85%) (Adeleke et al., 2015). The level of porosity of the soil was indicated by the presence of organic carbon (1.53%) while pH was confirmed to be 5.65 and the soil texture of the site was sandy-loam of which 76,15 and 9% were values for sand, clay and silt, respectively (Ganiyu et al., 2018). During the late planting season, a 3 × 5 factorial experiment fitted into Completely Randomized Design and Randomized Complete Block Design in both the screenhouse and the field, respectively. All experiments were set up with three replications. The experiment consisted of three tomato accessions, (CPTTO/19/191, CPTTO/19/193 and CPTTO/19/195). Titanium dioxide (TiO_2_) was prepared at four concentrations (0.3,0.7, 1.0, and 1.3 ml/l) while plots without TiO_2_ served as untreated control.

### Isolation and identification of *F. oxysporum* f. sp. *lycopersici*

Wilted tomato plants with yellow leaves were collected from wilt-endemic tomato field at Teaching and Research Farms of Federal University of Agriculture, Abeokuta and taken to the laboratory for fungal isolation. Tomato plant stems showing vascular discoloration were rinsed thoroughly in tap water and then macerated with a sterile scalpel. The macerated tomato plant stems were then surface sterilized using 1 % sodium hypochlorite, NaOCl for two (2) minutes, rinsed in three changes of sterile distilled water and dried on sterile filter paper. Segments from the stems were placed on Potato Dextrose Agar (PDA) medium amended with streptomycin sulphate (300 mg L_-1_) in petri-dishes and incubated at room temperature for 4 days. Sub-culturing of fungal isolates was done to obtain pure cultures of fungal isolate. Sub-culturing was done by single-spore isolation method on dried agar cultures. Identification of *F. oxysporum* f. sp. *lycopersici* was done under the light microscope and fungal structures were placed on slides, stained with methylene blue. Further identification using characteristic taxonomic and morphological features for *F. oxysporum* f. sp. *lycopersici* was conducted as contained in the work of (Leslie *et al*., 2006).

### Screenhouse experiment

#### Nursery and tomato transplanting

Three tomato seedlings were transplanted into each pot filled with 9 kg of steam-sterilized soil which was later thinned to one. There were forty–five (45) experimental pots altogether and each pot was arranged in rows of six with spacing of 0.5 m between the pots and 1 m between replicates.

#### Preparation and application of inoculum and TiO_2_

Conidia suspension of seven-day old pure cultures of isolated *F. oxysporum* f.sp. *lycopersici* were washed with sterile distilled water to obtain suspension of inoculums of the pathogen. The suspension was then filtered through one layer of Mira cloth, centrifuged, washed with sterile water and adjusted to a concentration of 10^6^ conidia per ml and was used to inoculate the four-week-old tomato seedlings at one (1) week after transplanting. Inoculation of the tomato seedlings was carried out using root-dip inoculation method at the rate of 1 ml / hole (Amini, 2009).

### Field experiment

#### Land preparation

The land was cleared, ploughed, harrowed. The experimental plot size was 3 m x 3 m with 1 m border row between plots were made. The experiment consisted of forty–five (45) experimental plots with thirty-six (36) tomato plants per plot.

#### Transplanting of tomato and application of TiO_2_

Tomato seedlings were transplanted at a spacing of 0.5 m x 0.5 m in the evening on the already prepared land. Tomato plants were treated with TiO_2_ at 2, 4 and 6 weeks after transplanting using soil drenching while plots TiO_2_ without served as control.

### Data collection and analysis

Data were collected on five sample plants at the middle of each experimental plot were plant height (cm), number of leaves/plant, number of flowers/plant, number of tomato fruits/plant, and fruit yield (tons/ha), disease incidence and disease severity. Disease assessment commenced at 4 and 6 weeks after transplanting.

Disease incidence was calculated by:

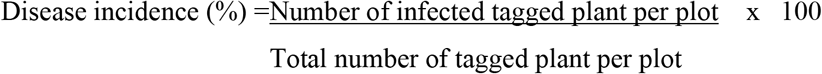

Severity was monitored through a visual scale (Lebeda and Buczkowski, 1986).

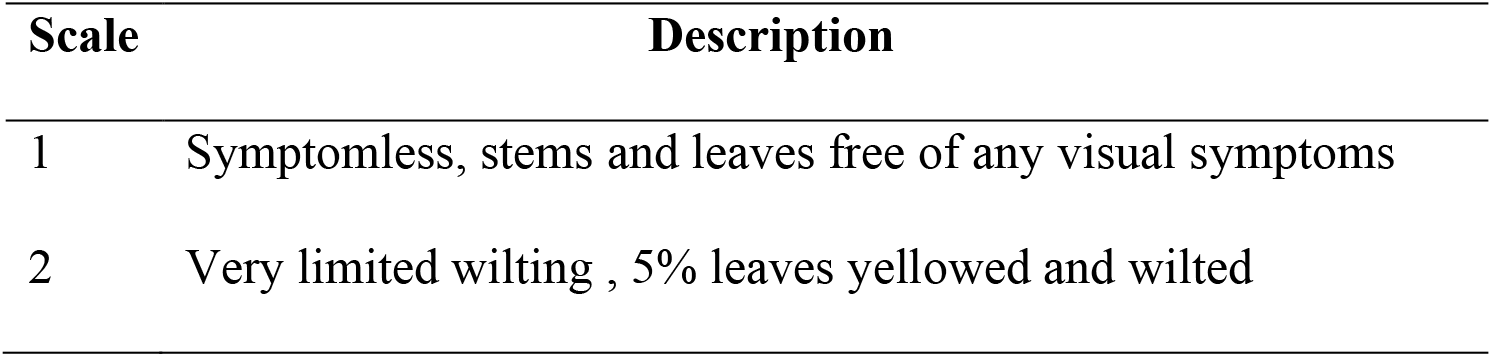
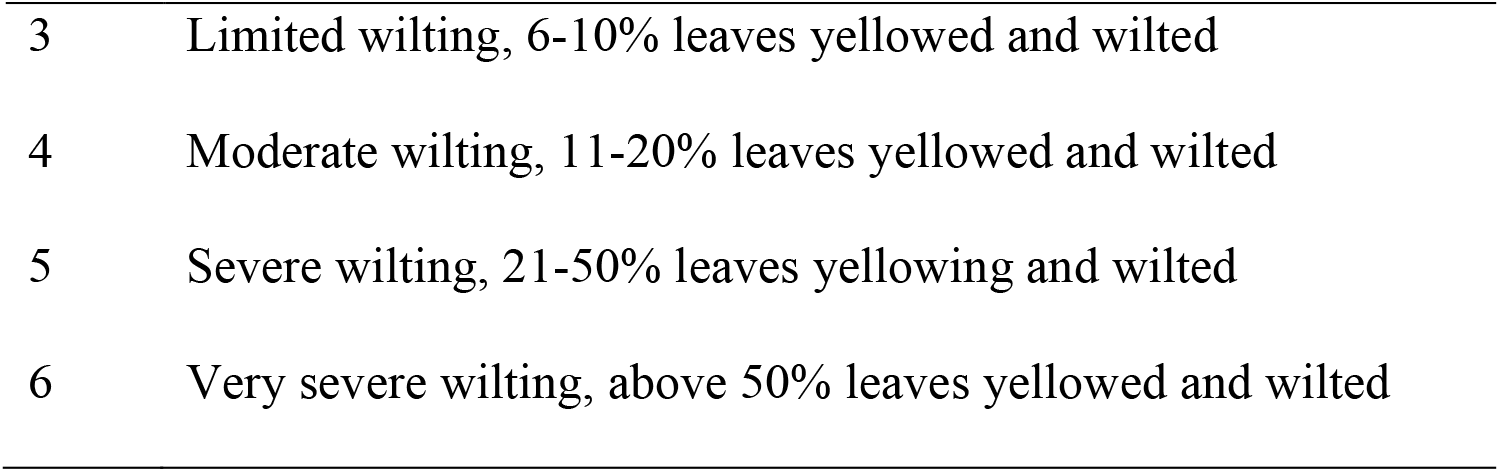

Data was subjected to analysis of variance (ANOVA) using Statistical Analysis System (SAS), 9.1 package and means were separated using the Duncan’s Multiple Range Test (p≤0.05).

## RESULTS

Table 1 shows the effect of TiO_2_ application on incidence of *Fusarium* wilt of tomato in both screenhouse and field experiments at four and six weeks after transplanting (WAT). Results from the screenhouse and field experiment indicated that the four treatments of TiO_2_ significantly reduced the incidence of *Fusarium* wilt in the three accessions of tomato compared to the control pots. However, treatment of TiO_2_ applied at 1.3ml/l concentration contributed to the lowest incidence of *Fusarium* wilt in both the screenhouse and field evaluation. In the screenhouse, the highest disease incidence of 52.00% and 82.20% were recorded in the untreated control pot containing CPTTO/19/193 and CPTTO/19/195 accessions, which were significantly different (p ≤ 0.05) from the lowest disease incidence of 2.30% and 10.10% recorded for CPTTO/19/191 treated with TiO_2_ at 1.3ml/l concentration at 4 and 6 WAT, respectively. On the field experiment, in untreated control plots at 4WAT, the highest disease incidence (60.00, 64.00 and 60.90%) were recorded for CPTTO/19/191, CPTTO/19/193 and CPTTO/19/195 accessions, respectively. These values were significantly higher than the lowest incidence of 3.50% recorded for CPTTO/19/191 treated with TiO_2_ at 1.3 ml/l concentration 4 WAT. Similarly, at 6 WAT the highest incidence (87.30, 92.80% and 90.10%) were recorded for CPTTO/19/191, CPTTO/19/193 and CPTTO/19/195 accessions, respectively, in the untreated control plots and were significantly different from the lowest incidence (20.70, 17.90 and 16.60%) recorded for each, treated with TiO_2_ at 1.3 ml/l concentration.

**Table 1:** Effect of TiO_2_ application on incidence of *Fusarium* wilt of tomato at four and six weeks after transplanting.

| Accession | $\text{TiO}_2$ (ml/l) | Screenhouse | | Field | |
| --- | --- | --- | --- | --- | --- |
|  |  | 4WAT<sup>†</sup> | 6 WAT | 4WAT | 6WAT |
| CPTTO/19/191 | 0.3 | 33.70<sup>bc</sup> | 62.90<sup>a-c</sup> | 43.30<sup>bc</sup> | 71.90<sup>a-c</sup> |
|  | 0.7 | 13.20<sup>bc</sup> | 43.50<sup>cd</sup> | 23.30<sup>ef</sup> | 55.50<sup>cd</sup> |
|  | 1.0 | 9.30<sup>f-h</sup> | 32.00<sup>de</sup> | 11.30<sup>f-h</sup> | 43.00<sup>de</sup> |
|  | 1.3 | 2.30<sup>h</sup> | 10.10<sup>f</sup> | 3.50<sup>h</sup> | 20.70<sup>f</sup> |
|  | Control | 50.00<sup>a</sup> | 77.50<sup>ab</sup> | 60.00<sup>a</sup> | 87.30<sup>ab</sup> |
| CPTTO/19/193 | 0.3 | 37.00<sup>b</sup> | 63.00<sup>a-c</sup> | 47.20<sup>b</sup> | 73.20<sup>a-c</sup> |
|  | 0.7 | 20.60<sup>c-e</sup> | 38.2<sup>de</sup> | 30.70<sup>c-e</sup> | 48.00<sup>de</sup> |
|  | 1.0 | 11.90<sup>e-g</sup> | 18.50<sup>ef</sup> | 18.90<sup>e-g</sup> | 28.80<sup>ef</sup> |
|  | 1.3 | 9.10<sup>f-h</sup> | 11.90<sup>f</sup> | 9.80<sup>f-h</sup> | 17.90<sup>f</sup> |
|  | Control | 52.00<sup>a</sup> | 81.50<sup>ab</sup> | 64.00<sup>a</sup> | 92.80<sup>a</sup> |
| CPTTO/19/195 | 0.3 | 22.20<sup>b-d</sup> | 64.60<sup>bc</sup> | 40.10<sup>b-d</sup> | 70.60<sup>bc</sup> |
|  | 0.7 | 18.30<sup>de</sup> | 38.40<sup>de</sup> | 28.10<sup>de</sup> | 48.60<sup>de</sup> |
|  | 1.0 | 8.60<sup>f-h</sup> | 12.30<sup>ef</sup> | 11.60<sup>f-h</sup> | 28.60<sup>ef</sup> |
|  | 1.3 | 4.10<sup>gh</sup> | 10.20<sup>f</sup> | 6.10<sup>gh</sup> | 16.60<sup>f</sup> |
|  | Control | 50.30<sup>a</sup> | 82.20<sup>ab</sup> | 60.90<sup>a</sup> | 90.10<sup>ab</sup> |

Means in the same column with different alphabet are significantly different ( $P \leq 0.05$ ) according to Duncan's Multiple Range Test, <sup>†</sup>WAT: Weeks After Transplanting

Table 2 shows the effect of TiO_2_ application on severity of *Fusarium* wilt of tomato in both screenhouse and field experiments at four and six weeks after transplanting. Application of TiO_2_ at 1.3 ml/l concentration showed significant reduction of *Fusarium* wilt severity in the three accessions of tomato used in both screenhouse and field experiments. In both the screenhouse and the field experiment, disease severity was generally at the peak on untreated control plants, ranged from 3.35-6.00 followed by 2.00-5.00 observed on tomato plants treated with TiO_2_ at 0.3 ml/l concentration. In screenhouse, lowest disease severity of 1.00 and 1.20 (CPTTO/19/191), 1.34 and 1.17 (CPTTO/19/193) and 1.00 and 1.63 (CPTTO/19/195) were recorded on plants treated with TiO_2_ at 1.3 ml/l concentration at 4 and 6WAT, respectively. These values were significantly different (p ≤ 0.05) from 4.43 and 5.00 (CPTTO/19/191), 3.35 and 4.43 (CPTTO/19/193) and 4.40 and 5.00 (CPTTO/19/195) obtained from untreated control plots. Similar trend was observed on the field. At 6WAT, the three accessions recorded highest disease severity (6.00) in untreated control plots while the lowest severity (1.67) was recorded in CPTTO/19/193 and CPTTO/19/195 treated with TiO_2_ at 1.3 ml/l concentration.

**Table 2:** Effect of TiO_2_ application on severity of *Fusarium* wilt of tomato plant at four and six weeks after transplanting.

| Accession | TiO<sub>2</sub> (ml/l) | Screenhouse |  | Field |  |
| --- | --- | --- | --- | --- | --- |
|  |  | 4WAT<sup>†</sup> | 6 WAT | 4WAT | 6WAT |
| CPTTO/19/191 | 0.3 | 2.33<sup>cd</sup> | 4.03<sup>a</sup> | 3.66<sup>cd</sup> | 5.00<sup>a</sup> |
|  | 0.7 | 2.20<sup>de</sup> | 3.33<sup>de</sup> | 3.00<sup>de</sup> | 4.00<sup>de</sup> |
|  | 1.0 | 1.03<sup>fg</sup> | 2.30<sup>f</sup> | 2.00<sup>fg</sup> | 3.00<sup>f</sup> |
|  | 1.3 | 1.00<sup>h</sup> | 1.20<sup>g</sup> | 1.30<sup>h</sup> | 2.00<sup>g</sup> |
|  | Control | 4.43<sup>a</sup> | 5.00<sup>a</sup> | 5.33<sup>a</sup> | 6.00<sup>a</sup> |
| CPTTO/19/193 | 0.3 | 2.00<sup>de</sup> | 3.53<sup>b-d</sup> | 3.00<sup>de</sup> | 4.33<sup>b-d</sup> |
|  | 0.7 | 2.34<sup>de</sup> | 2.34<sup>f</sup> | 3.00<sup>de</sup> | 3.33<sup>f</sup> |
|  | 1.0 | 1.35<sup>fg</sup> | 1.77<sup>f</sup> | 2.00<sup>fg</sup> | 2.67<sup>f</sup> |
|  | 1.3 | 1.34<sup>gh</sup> | 1.17<sup>g</sup> | 1.33<sup>gh</sup> | 1.67<sup>g</sup> |
|  | Control | 3.35<sup>bc</sup> | 4.43<sup>a</sup> | 4.33<sup>bc</sup> | 6.00<sup>a</sup> |
| CPTTO/19/195 | 0.3 | 3.45<sup>c</sup> | 4.00<sup>bc</sup> | 4.00<sup>c</sup> | 5.00<sup>bc</sup> |
|  | 0.7 | 2.24<sup>de</sup> | 3.24<sup>b-d</sup> | 3.00<sup>de</sup> | 4.33<sup>b-d</sup> |
|  | 1.0 | 1.33<sup>ef</sup> | 2.53<sup>ef</sup> | 2.33<sup>ef</sup> | 3.33<sup>ef</sup> |
|  | 1.3 | 1.00<sup>h</sup> | 1.63<sup>g</sup> | 1.00<sup>h</sup> | 1.67<sup>g</sup> |
|  | Control | 4.40<sup>ab</sup> | 5.00<sup>a</sup> | 5.00<sup>ab</sup> | 6.00<sup>a</sup> |

Means in the same column with different alphabet are significantly different ( $P \leq 0.05$ ) according to Duncan's Multiple Range Test, <sup>†</sup>WAT: Weeks After Transplanting

The effect of TiO_2_ on tomato plant indicated that the lowest value of plant height (3.40 cm, 3.43 cm and 3.44 cm) in the screenhouse at 2 WAT was recorded in the control pots for CPTTO/19/191, CPTTO/19/193 and CPTTO/19/195, respectively (Table 3). The highest plant height (19.33 cm) was recorded in pots containing CPTTO/19/195 treated with TiO_2_ at concentration of 0.7 ml/l.

**Table 3:** Effect of TiO_2_ application on plant height (cm) of tomato in the screenhouse and field test.

| Accession | TiO<sub>2</sub> (ml/l) | Screenhouse |  |  | Field layout |  |  |
| --- | --- | --- | --- | --- | --- | --- | --- |
|  |  | 2WAT<sup>†</sup> | 4WAT | 6WAT | 2WAT | 4WAT | 6WAT |
| CPTTO/19/191 | 0.3 | 8.67<sup>c</sup> | 18.43<sup>c-e</sup> | 29.03<sup>c-f</sup> | 10.67<sup>c</sup> | 22.67<sup>c-e</sup> | 33.00<sup>c-f</sup> |
|  | 0.7 | 10.43<sup>bc</sup> | 23.32<sup>a-d</sup> | 32.33<sup>a-f</sup> | 13.33<sup>bc</sup> | 27.83<sup>a-d</sup> | 36.50<sup>a-f</sup> |
|  | 1.0 | 13.37<sup>abc</sup> | 26.83<sup>ab</sup> | 36.34<sup>abc</sup> | 16.17<sup>abc</sup> | 30.83<sup>ab</sup> | 40.80<sup>abc</sup> |
|  | 1.3 | 15.86<sup>a</sup> | 30.53<sup>a</sup> | 46.32<sup>ab</sup> | 18.83<sup>a</sup> | 34.83<sup>a</sup> | 47.70<sup>ab</sup> |
|  | Control | 3.4<sup>d</sup> | 10.47<sup>fg</sup> | 16.23<sup>ef</sup> | 5.50<sup>d</sup> | 14.17<sup>g</sup> | 24.80<sup>ef</sup> |
| CPTTO/19/193 | 0.3 | 10.54<sup>c</sup> | 18.87<sup>de</sup> | 29.54<sup>cdef</sup> | 12.50<sup>c</sup> | 21.67<sup>de</sup> | 32.30<sup>cdef</sup> |
|  | 0.7 | 14.30<sup>abc</sup> | 21.21<sup>bcde</sup> | 33.45<sup>abcde</sup> | 16.00<sup>abc</sup> | 25.33<sup>bcde</sup> | 37.80<sup>abcde</sup> |
|  | 1.0 | 15.47<sup>ab</sup> | 25.23<sup>abc</sup> | 36.65<sup>abc</sup> | 18.67<sup>ab</sup> | 29.17<sup>abc</sup> | 40.80<sup>abc</sup> |
|  | 1.3 | 19.22<sup>a</sup> | 28.13<sup>ab</sup> | 44.87<sup>ab</sup> | 21.33<sup>a</sup> | 32.33<sup>ab</sup> | 47.00<sup>ab</sup> |
|  | Control | 3.43<sup>d</sup> | 9.65<sup>fg</sup> | 19.12<sup>f</sup> | 5.33<sup>d</sup> | 13.83<sup>fg</sup> | 23.70<sup>f</sup> |
| CPTTO/19/195 | 0.3 | 9.13<sup>c</sup> | 16.50<sup>ef</sup> | 27.24<sup>c-f</sup> | 11.17<sup>c</sup> | 20.50<sup>ef</sup> | 31.30<sup>c-f</sup> |
|  | 0.7 | 19.33<sup>c</sup> | 21.21<sup>b-e</sup> | 30.98<sup>b-f</sup> | 11.83<sup>c</sup> | 25.33<sup>b-e</sup> | 34.50<sup>b-f</sup> |
|  | 1.0 | 15.33<sup>ab</sup> | 27.20<sup>ab</sup> | 35.34<sup>a-d</sup> | 18.00<sup>ab</sup> | 30.00<sup>ab</sup> | 39.70<sup>a-d</sup> |
|  | 1.3 | 19.23<sup>a</sup> | 30.27<sup>a</sup> | 44.45<sup>a</sup> | 21.50<sup>a</sup> | 34.17<sup>a</sup> | 49.30<sup>a</sup> |
|  | Control | 3.44<sup>d</sup> | 10.23<sup>g</sup> | 22.43<sup>d-f</sup> | 5.67<sup>d</sup> | 13.5<sup>g</sup> | 26.00<sup>d-f</sup> |

Means in the same column with different alphabet are significantly different ( $P \leq 0.05$ ) according to Duncan's Multiple Range Test, <sup>†</sup>WAT: Weeks After Transplanting

CPTTO/19/191 treated with TiO_2_ at 1.3 ml/l concentration had the highest plant heights of 30.53 cm and 46.32 cm and were significantly different from the shortest plant heights (9.65 cm and 16.23 cm) observed in the control pot containing CPTTO/19/193 and CPTTO/19/191 at 4 and 6WAT, respectively. On the field, there were significant differences (p≤0.05) as the highest plant heights of 21.33 cm and 21.50 cm were recorded for CPTTO/19/193 and CPTTO/19/195 treated with TiO_2_ at concentration of 1.3 ml/l at 2WAT. Inversely, the shortest plant height of 5.50 cm, 5.33 cm and 5.67 cm were recorded for CPTTO/19/191, CPTTO/19/193 and CPTTO/19/195 accessions in the control plots. At 6WAT, there was significant difference (p≤0.05) as CPTTO/19/195 treated with TiO_2_ at concentration of 1.3 ml/l recorded significant higher plant height (49.30 cm) while the shortest plant height (23.70 cm) was recorded in the control plot with CPTTO/19/193.

Number of leaves per plant differed significantly (p ≤ 0.05) among the three accessions as shown in Table 4. At 2WAT, in screenhouse, the highest number of leaves (6.87) was recorded in CPTTO/19/193 treated with TiO_2_ at concentration of 1.00 ml/l while the lowest number of leaves (2.01) was recorded in CPTTO/19/195 treated with TiO_2_ at concentration of 0.3 ml/l. At the 4 and 6WAT, CPTTO/19/193 treated with TiO_2_ at concentration of 1.3 ml/l recorded the highest number of leaves (13.56 and 19.65), respectively while the lowest number of leaves (5.34) at 4WAT was recorded in the control pot of CPTTO/19/191. Also, control pot of CPTTO/19/193 recorded the lowest number of tomato leaves (10.54) at 6 WAT. On the field, the number of leaves 9.00 in CPTTO/19/193 when treated with TiO_2_ at concentration of 1.3 ml/l at 2WAT, was significantly higher than the lowest number of leaves 3.67 recorded in the control plot of CPTTO/19/195. At 4WAT, the three accessions, CPTTO/19/191, CPTTO/19/193 and CPTTO/19/195, recorded the highest number of leaves (16.00, 15.67 and 16.00), respectively, when treated with TiO_2_ at concentration of 1.3 ml/l. When treated with TiO_2_ at concentration of 1.3 ml/l, the number of leaves in CPT/19/193 (21.00) and CPTTO/19/195 (21.67) were significantly higher than the lowest number of leaves (12.33) recorded for CPTTO/19/191 in the control plots at 6WAT.

**Table 4:** Effect of TiO_2_ application on number of leaves of tomato plant in the screenhouse and field tests.

| Accession | TiO<sub>2</sub> (ml/l) | Screenhouse |  |  | Field |  |  |
| --- | --- | --- | --- | --- | --- | --- | --- |
|  |  | 2WAT<sup>†</sup> | 4WAT | 6WAT | 2WAT | 4WAT | 6WAT |
| CPTTO/19/191 | 0.3 | 3.34<sup>e-g</sup> | 6.32<sup>e-g</sup> | 11.23<sup>e-g</sup> | 5.00<sup>e-g</sup> | 9.00<sup>e-g</sup> | 14.00<sup>e-g</sup> |
|  | 0.7 | 3.43<sup>d-f</sup> | 7.39<sup>d-f</sup> | 13.30<sup>d-g</sup> | 5.33<sup>d-f</sup> | 10.33<sup>d-f</sup> | 15.00<sup>d-g</sup> |
|  | 1.0 | 3.47<sup>de</sup> | 10.12<sup>a-c</sup> | 15.78<sup>a-e</sup> | 5.67<sup>de</sup> | 13.67<sup>a-c</sup> | 18.00<sup>a-e</sup> |
|  | 1.3 | 5.54<sup>bc</sup> | 13.30<sup>a</sup> | 17.54<sup>a-c</sup> | 7.33<sup>bc</sup> | 16.00<sup>a</sup> | 20.67<sup>a-c</sup> |
|  | Control | 2.43<sup>f-i</sup> | 5.34<sup>fg</sup> | 19.34<sup>g</sup> | 4.00<sup>f-i</sup> | 8.00<sup>fg</sup> | 12.33<sup>g</sup> |
| CPTTO/19/193 | 0.3 | 4.03<sup>c-e</sup> | 7.43<sup>d-f</sup> | 11.45<sup>d-g</sup> | 6.00<sup>c-e</sup> | 10.33<sup>d-f</sup> | 14.67<sup>d-g</sup> |
|  | 0.7 | 4.67<sup>b-d</sup> | 9.45<sup>c-e</sup> | 12.54<sup>c-f</sup> | 6.67<sup>b-d</sup> | 11.33<sup>c-e</sup> | 16.67<sup>c-f</sup> |
|  | 1.0 | 6.87<sup>b</sup> | 10.23<sup>bc</sup> | 14.45<sup>a-e</sup> | 7.67<sup>b</sup> | 13.33<sup>bc</sup> | 18.00<sup>a-e</sup> |
|  | 1.3 | 6.43<sup>a</sup> | 13.56<sup>ab</sup> | 19.65<sup>ab</sup> | 9.00<sup>a</sup> | 15.67<sup>ab</sup> | 21.00<sup>ab</sup> |
|  | Control | 3.40<sup>e-g</sup> | 5.56<sup>e-g</sup> | 10.54<sup>fg</sup> | 5.00<sup>e-g</sup> | 8.67<sup>e-g</sup> | 13.00<sup>fg</sup> |
| CPTTO/19/195 | 0.3 | 2.01<sup>e-g</sup> | 7.05<sup>e-g</sup> | 13.87<sup>d-g</sup> | 5.00<sup>e-g</sup> | 9.00<sup>e-g</sup> | 15.67<sup>d-g</sup> |
|  | 0.7 | 3.98<sup>e-h</sup> | 7.45<sup>d-f</sup> | 12.54<sup>b-f</sup> | 5.00<sup>e-h</sup> | 10.67<sup>d-f</sup> | 17.00<sup>b-f</sup> |
|  | 1.0 | 4.31<sup>b-e</sup> | 9.73<sup>cd</sup> | 14.34<sup>a-d</sup> | 6.33<sup>b-e</sup> | 12.67<sup>cd</sup> | 18.67<sup>a-d</sup> |
|  | 1.3 | 5.43<sup>bc</sup> | 12.03<sup>a</sup> | 19.56<sup>a</sup> | 7.33<sup>bc</sup> | 16.00<sup>a</sup> | 21.67<sup>a</sup> |
|  | Control | 2.60<sup>i</sup> | 5.54<sup>g</sup> | 10.40<sup>fg</sup> | 3.67<sup>i</sup> | 7.00<sup>g</sup> | 13.00<sup>fg</sup> |

Means in the same column with different alphabet are significantly different ( $P \leq 0.05$ ) according to Duncan's Multiple Range Test, <sup>†</sup>WAT: Weeks After Transplanting

**Table 5:**
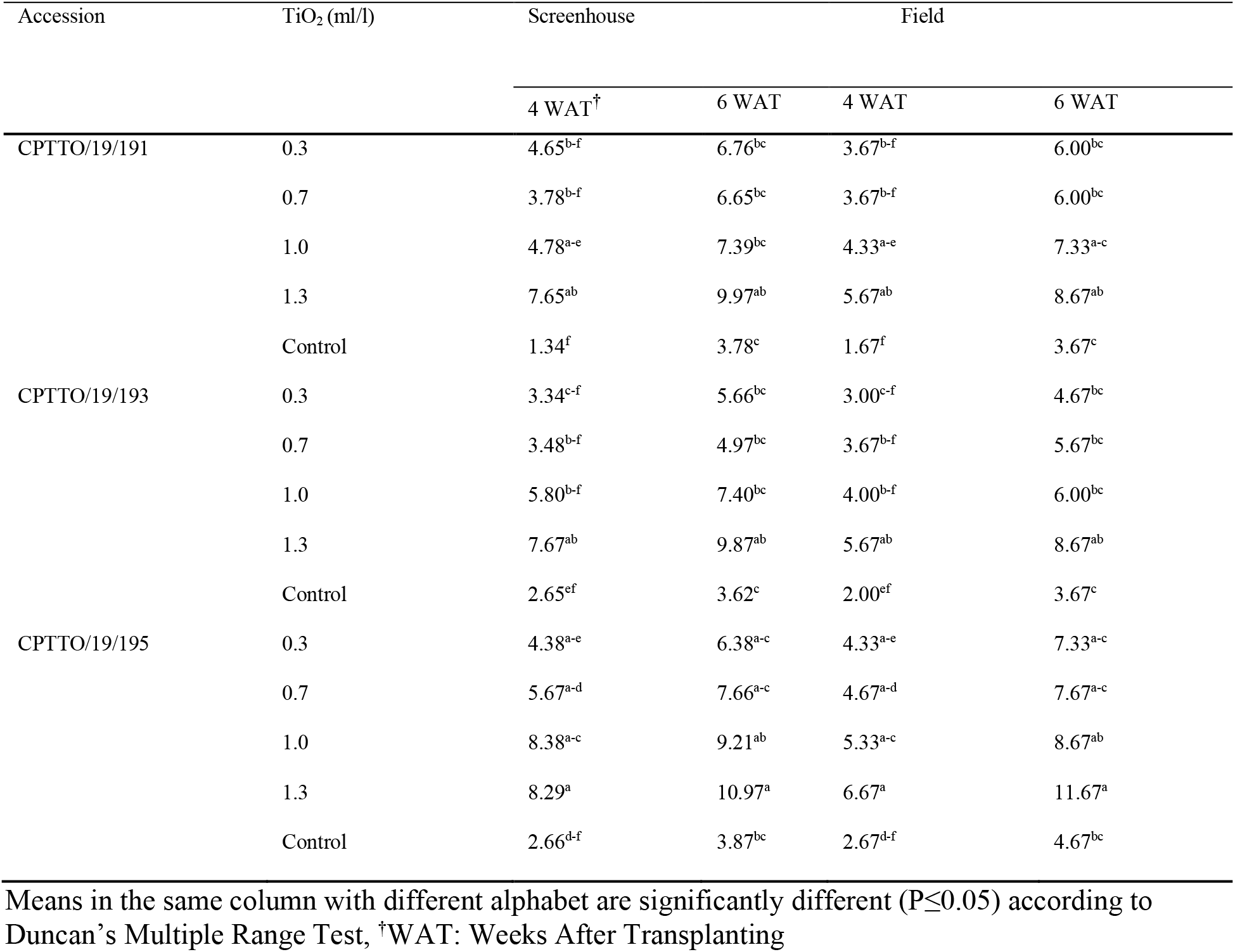
Effect of TiO_2_ application on number of flowers of tomato plant in the screenhouse and field tests.

At 4 WAT, CPTTO/19/195 treated with TiO_2_ at concentration of 1.3 ml/l recorded the highest number of flowers (8.38) which was significantly different from the lowest number of flowers (1.34) was recorded in control pot of CPTTO/19/191. The highest number of flowers (10.97) at 6 WAT was recorded in CPTTO/19/195 treated with TiO_2_ at concentration of 1.3 ml/l which was significantly different from the lowest number of flowers (3.62) was recorded in the control pot of CPTTO/19/193. On the field, highest number of flowers (6.67) was recorded in CPTTO/19/195 treated with TiO_2_ at concentration of 1.3 ml/l at 4 WAT which was significantly different from the lowest number of flowers (1.67) was recorded in the control plot of CPTTO/19/191. Furthermore, at 6 WAT, number of flowers (11.67) in CPTTO/19/195 treated with TiO_2_ at concentration of 1.3 ml/l was significantly higher than the number of flowers (3.67) recorded in the control plot of CPTTO/19/191 and CPTTO/19/193.

Table 6 shows the effect of TiO_2_ application on fruit yield (t/ha^-1^) of tomato. In screenhouse, CPTTO/19/193 treated with TiO_2_ at concentration of 1.3 ml/l had 2.74 t/ha^-1^ fruit yield, which was significantly higher than 0.59 t/ha^-1^ recorded for CPTTO/19/195 in the control pot. On the field, fruit yield of 28.00 t/ha^-1^ was recorded for CPTTO/19/193 treated with TiO_2_ at concentration of 1.3 ml/l which was significantly higher than fruit yields (5.30,4.50 and 6.70 t/ha^-1^) recorded for CPTTO/19/191, CPTTO/19/193 and CPTTO/19/195 in the control plots, respectively.

**Table 6:**
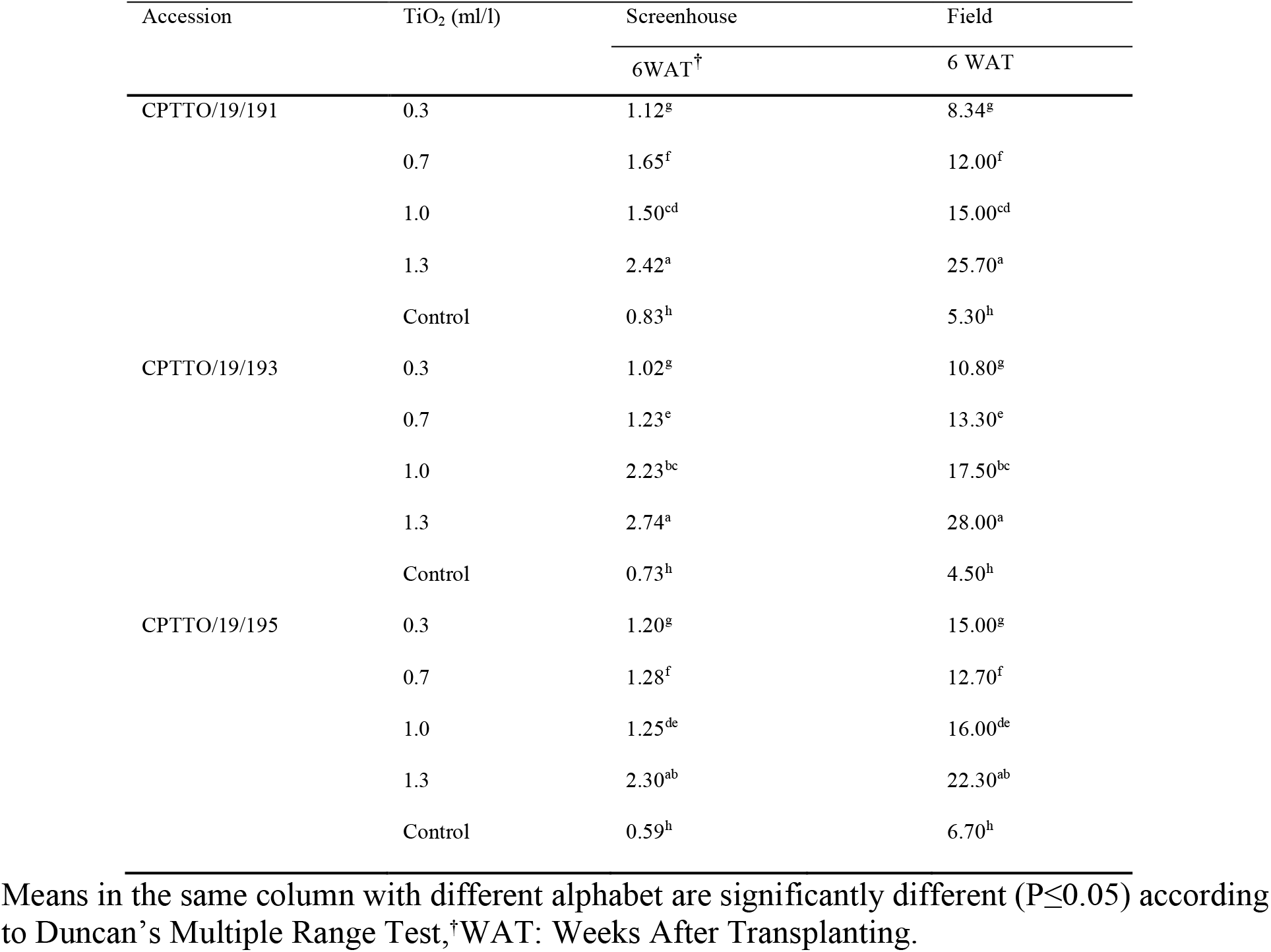
Effect of TiO_2_ application on fruit yield (t/ha^-1^) of tomato.

## DISCUSSION

The results obtained showed that application of TiO_2_ at higher concentration of 1.3 m/l significantly reduced the incidence and severity of *Fusarium* wilt of the tomato accessions compared to the other level of treatment application. This confirmed earlier result that demonstrated that TiO_2_ were potent on soil borne fungi (Frazer, 2001). TiO_2_ has also been shown to be effective in controlling *Fusarium* wilt of Basil (*Ocimum basilicum*) caused by (*Fusarium oxysporum* f.sp. *basilici*) (Adams *et al*., 2003). Application of TiO_2_ at lower concentration may not show only the right direction impact but also caused a positive impact on plant growth and control of fungal and bacteria diseases.

Disease incidence and disease severity were significantly reduced with corresponding increase in the number of leaves, plant height, number of flower and yield by the application of TiO_2_ at higher concentrations. The application of TiO_2_ at concentration of 1.3 ml/l had the best positive effect on tomato. The higher the concentration the higher the yield in the three tomato accessions treated with TiO_2_ while a lower yield and other parameters were recorded in the control treatment. Unlike on disease incidence and severity, the higher the concentration the lower the disease exhibited by the plant treated with TiO_2._ The control plant showed the highest disease. This result is in accordance with what was reported by (Chao *et al*., 2005), that the application of titanium dioxide (TiO_2_) on food crops promote plant growth, increase the photosynthetic rate, reduce disease severity and enhance yield by 30%. Titanium dioxide (TiO_2_) has also been shown to possess a strong oxidation reaction, which can target organic compounds. It has also been shown to be very effective in inactivating avian influenza virus (Cui *et al*., 2010). Titanium dioxide (TiO_2_) has also been recommended for control of other plant diseases (Cui *et al*., 2009).

*Fusarium* wilt disease of tomato is economically important infection to the crop worldwide. *Fusarium* wilt is controlled through many strategies such as usage of resistant varieties, biological control, host defense induction and integrated management. The use of titanium dioxide has been proven to be a useful tool for induced resistance studies in tomato as observed in this study.

## CONCLUSION

The overall result of this research work provided a basis for a major conclusion. Higher concentration of titanium dioxide application (TiO_2_) at concentrations of 1.3 ml/l had significant effect on *Fusarium* wilt of tomato; it reduced the incidence and severity of *Fusarium* wilt of tomato. Also, its application resulted in a significant increase in the yield of tomato. The protection of plants against pathogen using titanium dioxide is a promising control strategy. It can become an important component of pest management programs, particularly in cases where current control measures are less effective. Obviously, one of the outcomes of the use of titanium dioxide should be a reduction of the use of fungicides which is of major concern in the preservation of environment. Therefore, application of TiO_2_ at 1.3 m/l for the management of *Fusarium* wilt of tomato is feasible.

## Notes

### Competing Interest Statement

The authors have declared no competing interest.

